# A fast and efficient strategy for the NMR assignment of Fab methyl groups

**DOI:** 10.1101/2025.10.21.682867

**Authors:** Faustine Henot, Béatrice Vibert, Arthur Giraud, Sarra Dbira, Lionel Imbert, Adrien Favier, Peter Güntert, Séverine Clavier, Elodie Crublet, Camille Doyen, Jérôme Boisbouvier, Oriane Frances

## Abstract

Owing to their high specificity and therapeutic effectiveness, monoclonal antibodies (mAbs) have rapidly become one of the leading classes of biologic drugs used to treat critical illnesses. The antigen-binding fragment (Fab) of mAbs plays a key role in the antigen recognition, so its structural characterization is essential, as even a slight change to its Higher Order Structure (HOS) can impact the antibody’s potency. Recently, 2D methyl NMR has been introduced as a powerful method to assess both the structure and integrity of therapeutic Fab fragments. However, the identification of methyl group resonances in NMR spectra remains rare since Fabs are large heterodimers of ∼50 kDa. Here, we present the methyl group assignment of an IgG1 Fab produced in a cell-free system with an optimal isotope labelling. We first assigned 99% of the alanine, isoleucine, leucine, methionine, and valine methyl groups of the therapeutic Fab targeting LAMP1 antigen. Building on this assignment, we propose a “divide and conquer” strategy that exploits sequence identities to rapidly assign methyl groups of other IgG1 Fabs. We demonstrate that the assignment of the Fab’s constant region can easily be transferred from one IgG1 to another and that the variable part of a new Fab can be assigned using smaller uniformly ^15^N,^13^C-labelled constructs. We applied our strategy to ipilimumab’s Fab and, using the assignment of ipilimumab’s variable part and Fab anti-LAMP1’s constant part, we could transfer the assignment of 89% of the methyl-containing amino acids to the entire ipilimumab Fab without having to produce deuterated samples. This assignment strategy can be generalised to any other IgG1 Fabs provided that their constant regions are identical and the strategy can be adapted to accommodate the expression levels of the different variable domains. This new method drastically facilitates the Fab assignment process, making it suitable for the pharmaceutical timeline.

## Introduction

Monoclonal antibody (mAb) production in pharmaceutical companies is subject to strict specifications especially regarding product purity, stability, and structural characterization to ensure patients’ safety. Among these, the structure of a biological product, being directly linked to its function, is of paramount importance. Chemical modifications possibly occuring during mAb production or arising from product long-term storage can potentially affect the bioproduct’s structure and alter its potency. While conventional mass spectrometry analysis can inform about chemical modifications of the primary structure, it does not allow to link potency changes observed during biological activity tests with structural changes. Orthogonal biophysical tools currently available in the pharmaceutical industry, such as intrinsic fluorescence, differential scanning calorimetry or circular dichroism, lack sufficient resolution to highlight subtle changes in mAbs’ structures that have an impact on their potency. To solve this issue and bridge the gap between biological activity changes and biological product structural modifications, NMR methods have been introduced. Using NMR, it is possible to record well resolved spectral fingerprints of mAbs at natural abundance (Brinson et al. 2018; Marino et al. 2015) that allow to monitor structural modifications induced by stress tests performed on mAbs (Cerofolini et al. 2023). However, to investigate structural changes at atomic resolution it is necessary to assign each detected NMR signal to the corresponding atom of the mAb and to locate them in the structure. Several studies have described different protocols either using yeast, bacterial, mammalian or cell-free expression systems to produce labelled mAb fragments and assign their backbone resonances (Liu et al. 2007a, Liu et al. 2007b, Yagi et al. 2015, Ghasriani et al. 2022, Solomon et al. 2023, Gagné et al. 2024, Giraud et al. 2024, Sarker and Aubin 2024). Nonetheless, for pharmaceutical quality control studies, methyl group resonances present various advantages over backbone resonances. Indeed, the proton multiplicity, the greater natural abundance of ^13^C over ^15^N (1.1% versus 0.37%) and the favorable relaxation properties of the methyl groups make them particularly suitable probes for NMR fingerprint comparisons. Despite its relevance, methyl group resonance assignment of mAb fragments remains scarce in the literature. Previous attempts allowed the assignment of methyl groups of both a homodimeric 26 kDa reduced human IgG1 CH_3_ domain (Liu et al. 2007a) and a 26 kDa chimeric ScFv construct (Ghasriani et al. 2022). Fab fragments being larger heterodimers (∼50 kDa), their methyl group NMR assignment is more complex and requires the production of ^2^H, ^15^N, ^13^C labelled samples. Gagné et al. and Sarker and Aubin recently presented the first isoleucine, leucine, and valine methyl group assignments of two Fab fragments produced in *E. coli* (Gagné et al. 2024, Sarker and Aubin 2024).

In this article we present for the first time the nearly complete assignment of alanine, isoleucine, leucine, valine, methionine and threonine methyl groups of a therapeutic Fab fragment. This challenging task was performed on the Fab fragment of anti-LAMP1 IgG1 produced in a cell-free system with optimal perdeuteration and methyl specific labelling. Anti-LAMP1 mAb was selected as it interacts with the lysosomal-associated membrane protein 1 (LAMP1); a glycosylated protein primarily located on the lysosomal membrane that is expressed at the surface of tumor cells (Sarafian et al. 1998, Terasawa et al. 2016) and has been associated with tumor development and metastatic progression (Agarwal et al. 2014, Alessandrini et al. 2017, Cameron et al. 2020). Using this assignment of anti-LAMP1 Fab as a starting point, we developed an efficient method to assign other Fab fragments based on the “divide and conquer” approach and exploiting sequence identities between Fabs. Such an approach simplifies sample preparation, as it does not require complex preparation of expensive perdeuterated samples, and is compatible with the pharmaceutical timeline.. A Fab consists of a light and a heavy chain, each composed of a variable domain and a highly conserved constant domain. Our strategy relies on sequence identity between targeted IgG1 Fab fragments. We demonstrate that already made assignments of the constant part of anti-LAMP1 Fab can be transferred to the constant part of a new IgG1 Fab fragment: ipilimumab. The variable domain of ipilimumab could then be assigned with a minimal number of uniformly ^15^N,^13^C-labelled small constructs. We show that combining the assignment of ipilimumab’s variable part together with that of the already assigned anti-LAMP1’s constant part yields almost complete methyl assignments for the full ipilimumab Fab without large scale production of deuterated Fab samples. Our method significantly accelerates Fab assignment, a defining prerequisite to monitor higher order structure changes by NMR, and provides new opportunities to shorten the research-to-market delay.

## Materials and methods

### Protein plasmid preparations

For anti-LAMP1 Fab constructs, the light chain (LC - residues D1 to C213) and heavy chain (HC – residues Q1 to T226) sequences of the anti-LAMP1 mAb (Pruvost et al. 2023, Giraud et al. 2024) were cloned separately in pIVEX 2.4d and pIVEX 2.3d plasmids between NotI and SmaI and between NcoI and SmaI, respectively. Each plasmid contained either an N-terminal 6-His tag (LC) or a Twin-Streptag (HC) followed by a TEV protease cleavage site (see supplementary information for detailed sequences).

For ipilimumab constructs, the light chain (LC - residues E1 to C215) sequence (Lee et al. 2019) was cloned in pIVEX 2.3d plasmid between NcoI and SmaI. The heavy chain (HC – residues Q1 to K223), the Single chain Fragment variable (ScFv – residues Q1 to S118 from the heavy chain linked to residues E1 to T110 of the light chain with a linker composed of four repeats of the sequence GGGGS (Huston et al. 1988, Ghasriani et al. 2022)), the variable part of the light chain (VL – residues E1 to T110) and the variable part of the heavy chain (VH – residues Q1 to S118) sequences of ipilimumab were cloned separately in pIVEX 2.4d plasmids between NotI and SmaI. Each ipilimumab construct, except the LC, contained an N-terminal 6-His-tag followed by a TEV protease cleavage site (ENLYFQG). A glycine residue was added either before the E1 or the Q1 to initiate translation with a small amino acid. (see supplementary information for detailed sequences).

Each of the seven different plasmids was transformed into *E. coli* strain TOP10 grown in LB medium (500 mL) with 100 μg/mL of ampicillin. Plasmids were then purified using the NucleoBond Xtra Maxi Plus kit (Macherey-Nagel) and the final elution concentration was adjusted to approximatively 1 mg/mL of DNA in RNase-free water.

### In-vitro samples optimization and production

The S30 lysates were prepared based on a protocol previously described (Imbert et al. 2021) using an *E. coli BL21 (DE3)* strain. DTT was omitted in the S30 buffer dialysis batches to prevent a possible interaction with disulfide bridges of the Fab, ScFv, VL and VH fragments. All reactions were performed at 30 °C with gentle shaking (20 rpm) using a hybridization oven (Techne) in 1.5 mL Eppendorf tubes for small-scale syntheses and in 3 mL GeBAflex dialysis devices (Euromedex) for larger-scale syntheses.

For optimization of betaine, magnesium and DsbC concentrations, small-scale cell-free syntheses of the Fab fragments, ScFv, VL, and VH constructs were undertaken individually in 50 μL batch mode. Reactions were stopped after 3 h and analyzed with Western blots using an anti-His-tag or anti-Fab antibody. The best conditions are summarized in supplementary table S1.

For larger reaction volumes, the continuous exchange cell-free (CECF) mode was used with a reaction volume of 3 mL and a reaction-to-dialysis volume ratio of 1:5. For the reaction mixture and the dialysis mixture, standard cell-free components (nucleotides, HEPES, folinic acid, cyclic AMP, spermidine, NH_4_OAc and creatine phosphate) were added as described in a previously published protocol (Imbert et al. 2021). Betaine and magnesium were added at concentrations described in table S1. To form disulfide bridges, both reduced and oxidized glutathione were used at concentrations of 0.9 and 3.6 mM, respectively, according to previously published protocols (Yin and Swartz 2004, Giraud et al. 2024). The 20 amino acids were added as a mix at a final concentration of 1 mM each. The labelling scheme used for each sample is presented in tables S2 and S3. To produce uniformly ^13^C, ^15^N labelled samples, the amino acid mixture was replaced with an algal-derived amino acid mix (Isogro® ^13^C, ^15^N powder growth medium) hydrolysed as described previously (Imbert et al. 2021), supplemented with unlabelled tryptophan and cysteine at a final concentration of 1 mM each. Samples specifically labelled on methyl groups were obtained by adding each of the 20 amino acids with the desired labelling directly to the cell-free reaction mixture. For Ile-δ_1_, Leu-δ*-pro-S/pro-R*, and Val-γ*-pro-S/pro-R*, labelled precursors were added instead of the corresponding amino acids. 2-keto-3-(CD_3_-methyl)-3,4-(D_2_)-5-(^13^C)-pentanoate was used to selectively label isoleucine at the δ_1_ position. α-ketoisovaleric acid 3-D_1_, containing two ^13^CH_3_ methyl groups, was used to label valine at both γ*-pro-R* and γ*-pro-S* positions simultaneously. A variant with one ^13^CH_3_ and one ^12^CD_2_ methyl group was employed to achieve 50% labeling at γ*-pro-R* and 50% at γ*-pro-S*, without stereospecificity. Uniformly ^13^C-labeled α-ketoisocaproic acid was used to label leucine at both δ*-pro-R* and δ*-pro-S* positions.

Finally creatine kinase, T7 RNA polymerase, tRNA, DsbC, S30 *E. coli BL21(DE3)* bacterial extract and the DNA plasmids were added to the reaction mixture. For ScFv, VL, and VH constructs, a single plasmid was used for their production. All plasmids were added at a total concentration of 16 μg/mL in CECF mode. For the Fab constructs, both chains were co-expressed *in vitro*. LC:HC ratios of 3:1 and 1:2.2 were found to be optimal for ipilimumab and anti-LAMP1 production, respectively. Each 3 mL GeBAflex was placed in a 25 mL centrifugation tube containing 15 mL of dialysis mixture. CECF reactions were stopped after 16 h and mAb fragments were purified using the purification protocols described below.

### Ipilimumab and anti-LAMP1 fragments’ purification

CECF reaction mixtures were centrifuged for 10 min at 14,000 rpm. Supernatants were diluted by a factor of 4 either in 150 mM NaCl, 20 mM Na_2_HPO_4_/NaH_2_PO_4_ at pH 7.4 (for both Fabs, ScFv and VL fragments) or in 2X PBS, 20 mM imidazole at pH 7.4 (for the VH fragment). The diluted supernatants were purified using affinity chromatography columns. A HiTrap protein L column of 1 mL was used for both Fabs, ScFv and VL fragments whilst a HisTrap FF column of 1 mL was used for VH fragment purification. (See supplementary table S4 for detailed purification protocols and buffer compositions).

VL, ScFv and Fab fragments were all buffer-exchanged and concentrated in the NMR buffer containing 100% D_2_O, 50 mM MES at pH 6.9, 100 mM NaCl and cOmplete® protease inhibitors using a centrifugal filtration unit (Amicon® 10 kDa molecular weight cutoff (MWCO) for ScFv and Fab fragments, or 3 kDa MWCO for the VL fragment). The final samples were placed in either 4 mm Shigemi NMR tubes (for uniformly labelled and highly concentrated samples) or in 3 mm NMR tubes for other samples (see tables S2 and S3 for details on the samples).

### NMR Spectroscopy

For the anti-LAMP1 Fab fragment, all NMR experiments were recorded at 308 K on a Bruker Avance III HD spectrometer equipped with a cryogenic probe operating at ^1^H frequency of 950 MHz. 2D ^1^H-^13^C SOFAST methyl TROSY (Amero et al. 2009) experiments were recorded to identify each methyl type. For residues containing two methyl groups, only one was selectively labeled to facilitate spectral analysis (see supplementary table S2 for details about samples’ labelling). Connection of isoleucines, alanines, and valines-γ-*pro-R* methyl signals to previously assigned backbone frequencies (Giraud et al. 2024) was undertaken using 3D “out and back” HCC, HC(C)C and HC(CC)C experiments (Tugarinov and Kay 2003) acquired using a 0.26 mM sample of U-[^2^H, ^12^C], Ala-[2-^2^H; 1,2-^13^C_2_; [^13^C^1^H_3_]^β^], Ile-[2,3,4,4-^2^H_4_; 1,2,3,4-^13^C_4_; [^13^C^1^H_3_]^δ1^/[^12^C^2^H_3_]^γ2^], Val-[2,3-^2^H_2_; 1,2,3-^13^C_3_; [^13^C^1^H_3_]^*pro-R*^/[^12^C^2^H_3_]^*pro-S*^] anti-LAMP1 Fab fragment. A 3D CCH HMQC-NOESY-HMQC NMR experiment (Tugarinov et al. 2005, Törner et al. 2020) was acquired using a 0.31 mM sample of U-[^2^H, ^12^C], Ala-[^13^C^1^H_3_]^β^, Ile-[^13^C^1^H_3_]^δ1^, Val-[^13^C^1^H_3_]^*pro-R*^, Thr-[^13^C^1^H_3_]^γ^, Met-[^13^C^1^H_3_]^ε^, Leu-[^13^C^1^H_3_]^*pro-S*^ anti-LAMP1 Fab fragment. The NOE mixing time was set to 275 ms.

For ipilimumab fragments, NMR experiments were recorded at 308 K except for the ScFv fragment, for which NMR experiments were run at 313 K, using Bruker Avance III HD spectrometers equipped with cryogenic probes. 2D ^1^H-^13^C SOFAST methyl TROSY (Amero et al. 2009) experiments were recorded to identify each methyl type for the VL, ScFv, and Fab fragments of ipilimumab (see table S3 for details about samples). For backbone sequential assignment, two sets of 3D triple-resonance experiments (Favier and Brutscher 2019) were acquired on Bruker Avance III HD spectrometers equipped with cryogenic probes using either a 0.37 mM sample of U-[^15^N, ^13^C] VL fragment or a 0.22 mM sample of U-[^15^N, ^13^C] ScFv fragment (supplementary table S3). Methyl group assignment of the VL and ScFv fragments of ipilimumab was undertaken using 3D hCCH TOCSY experiments with transfer delays of 10 to 20 ms, the interscan delay was set at 1.3 s, the acquisition times were adjusted to either 7.5 or 11 ms in the ^13^C indirect dimensions, and to 76 ms in ^1^H direct dimension. Additionally, a 3D HCH-NOESY experiment was recorded with an interscan delay set to 1.0 s. The acquisition times in the ^13^C and ^1^H indirect dimensions were set to 11-12 ms (*t*_1max_) and to 19-20 ms (*t*_2max_). In the ^1^H direct dimension, *t*_3max_ was fixed to 50 ms. The NOE mixing period was set to 120-150 ms.

All data were processed and analyzed using nmrPipe/nmrDraw (Delaglio et al. 1995), Topspin 4.1.4 (Bruker), CcpNmr (Vranken et al. 2005) and MethylFLYA (Pritišanac et al. 2019).

## Results and discussion

### NMR assignment of anti-LAMP1 Fab methyl group resonances

To reach the goal of enabling a fast and efficient assignment of Fab fragments’ methyl groups using NMR, the assignement of methyl groups of a first Fab fragment was required. Several methods, such as transfer from the backbone assignment to methyl groups and structure-based approaches using NOESY experiments, have already been widely described and frequently used to assign methyl group resonances of complex targets including dynamic proteins and large molecular assemblies (Tugarinov and Kay 2003, Ayala et al. 2009, Mas et al. 2013, Xu and Matthews 2013, Chao et al. 2014, Pritišanac et al. 2017, Monneau et al. 2017, Pritišanac et al. 2019, Nerli et al. 2021, Henot et al. 2021, Törner et al. 2021). Here, we combined these two methods to assign the Fab fragment of anti-LAMP1.

Backbone sequential assignment of the anti-LAMP1 Fab had already been undertaken, leading to 89% of ^1^H^N^, 89% of ^15^N^H^, 92% of ^13^C’, 93% of ^13^C^α^ and 89% of ^13^C^β^ resonances assigned (Giraud et al. 2024). “Out and back” HCC, HC(C)C and HC(CC)C experiments enabled to connect previously assigned backbone C^α^ and C^β^ resonances to alanine-β, valine-γ-*pro-R*, and isoleucine-δ_1_ methyl signals. 53% of alanine-β, 43% of valine-γ-*pro-R*, and 33% of isoleucine-δ_1_ could be unambiguously assigned (Fig. 1). These assigned methyl group resonances, together with the amino acid type of each methyl signal, the 3D structure of the anti-LAMP1 Fab fragment (Pruvost et al. 2023) and the 434 intermethyl NOE cross peaks recorded on the U-[^2^H, ^14^N, ^12^C], Ala-[^13^C^1^H_3_]^β^, Ile-[^13^C^1^H_3_]^δ1^, Val-[^13^C^1^H_3_]^*pro-R*^, Thr-[^13^C^1^H_3_]^γ^, Met-[^13^C^1^H_3_]^ξ^, Leu-[^13^C^1^H_3_]^*pro-S*^ anti-LAMP1 Fab fragment were given as input to the software MethylFLYA (Pritišanac et al. 2019) (Fig. 1).

**Fig. 1.**
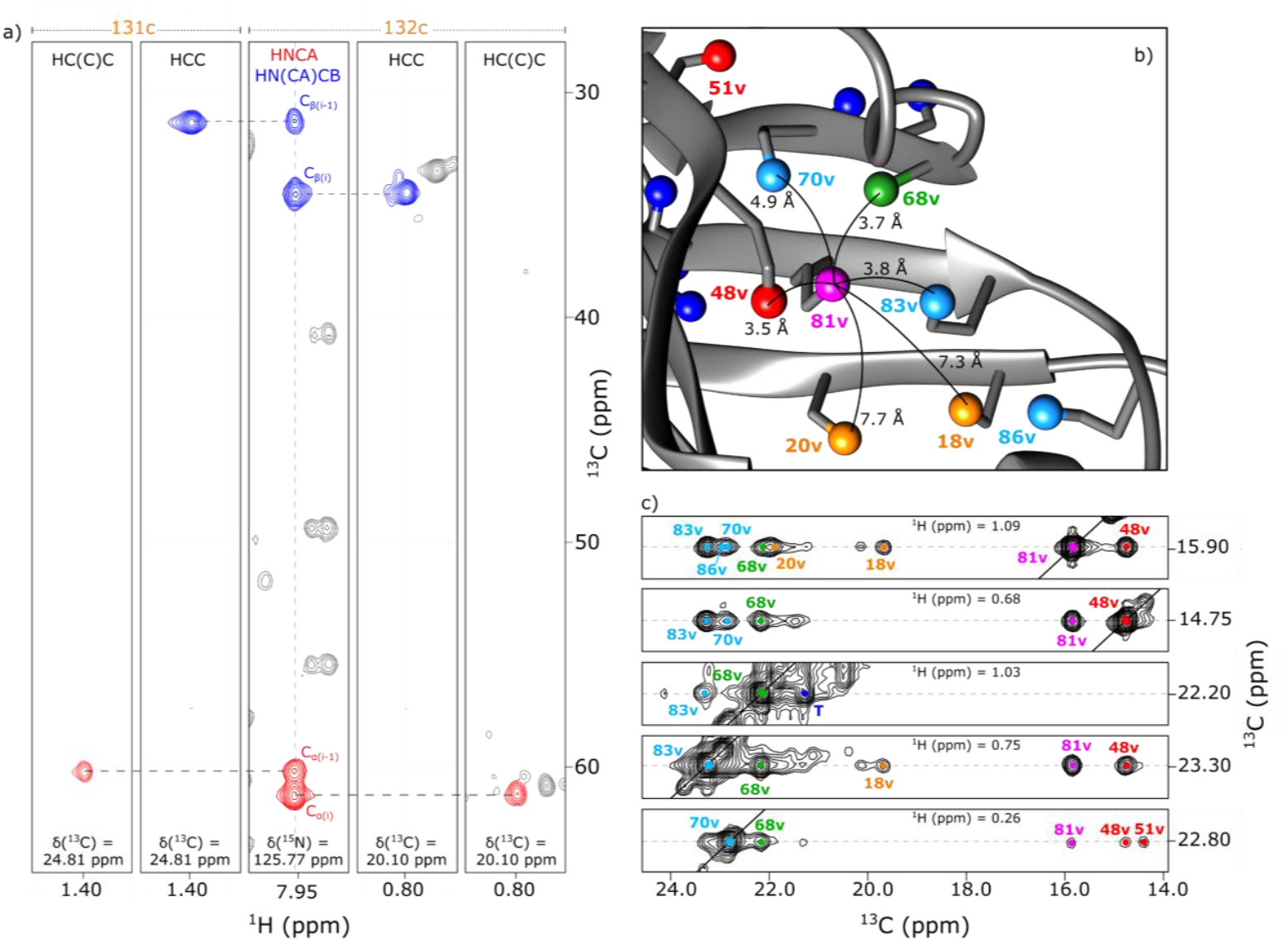
Methyl group assignment strategy for anti-LAMP1 Fab. **a)** Assignment transfer from the backbone to Val^*pro-R*^ methyl groups. 2D-extracts from 3D ‘out and back’ HCC and HC(C)C experiments correlating methyl resonances with ^13^C^β^ (blue) and ^13^C^α^ (red) backbone resonances are displayed. An anti-LAMP1’s Fab fragment labelled U-[^2^H, ^12^C], Ala-[2-^2^H; 1,2-^13^C_2_; [^13^C^1^H_3_]^β^], Ile-[2,3,4,4-^2^H_4_; 1,2,3,4-^13^C_4_; [^13^C^1^H_3_]^δ1^/[^12^C^2^H_3_]^γ2^], Val-[2,3-^2^H_2_; 1,2,3-^13^C_3_; [^13^C^1^H_3_]^*pro-R*^/[^12^C^2^H_3_]^*pro-S*^] was used to record these experiments. The middle panel shows a 2D extract of HNCA and HN(CA)CB for Val-132c allowing to connect methyl groups of Val-132^*pro-R*^ and Val-131^*pro-R*^ to the previously assigned backbone resonances. **b)** Zoom extracted from the 3D structure of anti-LAMP1 Fab and focusing on the variable part of the heavy chain (PDB 8ATH, Pruvost et al. 2023). The zoom shows methyl groups involved on the traces shown in panel c. The methyl groups are represented by spheres. Alanines-β, isoleucines-δ_1_, valines-*pro-R*, leucines-*pro-S*, threonines-γ, and methionines-ε are depicted in green, red, orange, light blue, dark blue, and purple, respectively. **c)** 2D-extracts from a 3D HMQC-NOESY-HMQC using an anti-LAMP1’s Fab fragment sample U-[^2^H] and specifically labelled on Ala-[^13^C^1^H_3_]^β^, Met-[^13^C^1^H_3_]^ε^, Leu-[^13^C^1^H_3_]^*pro-S*^, Val-[^13^C^1^H_3_]^*pro-R*^, Ile-[^13^C^1^H_3_]^δ1^, and Thr-[^13^C^1^H_3_]^γ^. Planes were extracted at the methyl proton frequencies of M81v, I48v, A68v, L83v and L70v

The automated methyl group assignment software was run with distance cutoffs of 5.5, 6.0 and 6.5 Å and 100 runs per distance cutoff. 10 additional A / I / V residues (7% of methyl groups) for which ambiguous assignment was obtained using “out and back” HCC experiments were assigned with high confidence using MethylFLYA. With this additional input, MethylFLYA was used to complete the assignment of alanines, valines, isoleucines, methionines, leucines, and threonines. Combining iterative MethylFLYA runs with a careful and intensive manual analysis of the NOESY spectrum, 96% of leucine-δ-*pro-S* and 100% of isoleucine-δ_1_, alanine-β, valines-γ-*pro-R* and methionines-ε methyls were assigned (Fig. 2).

**Fig. 2.**
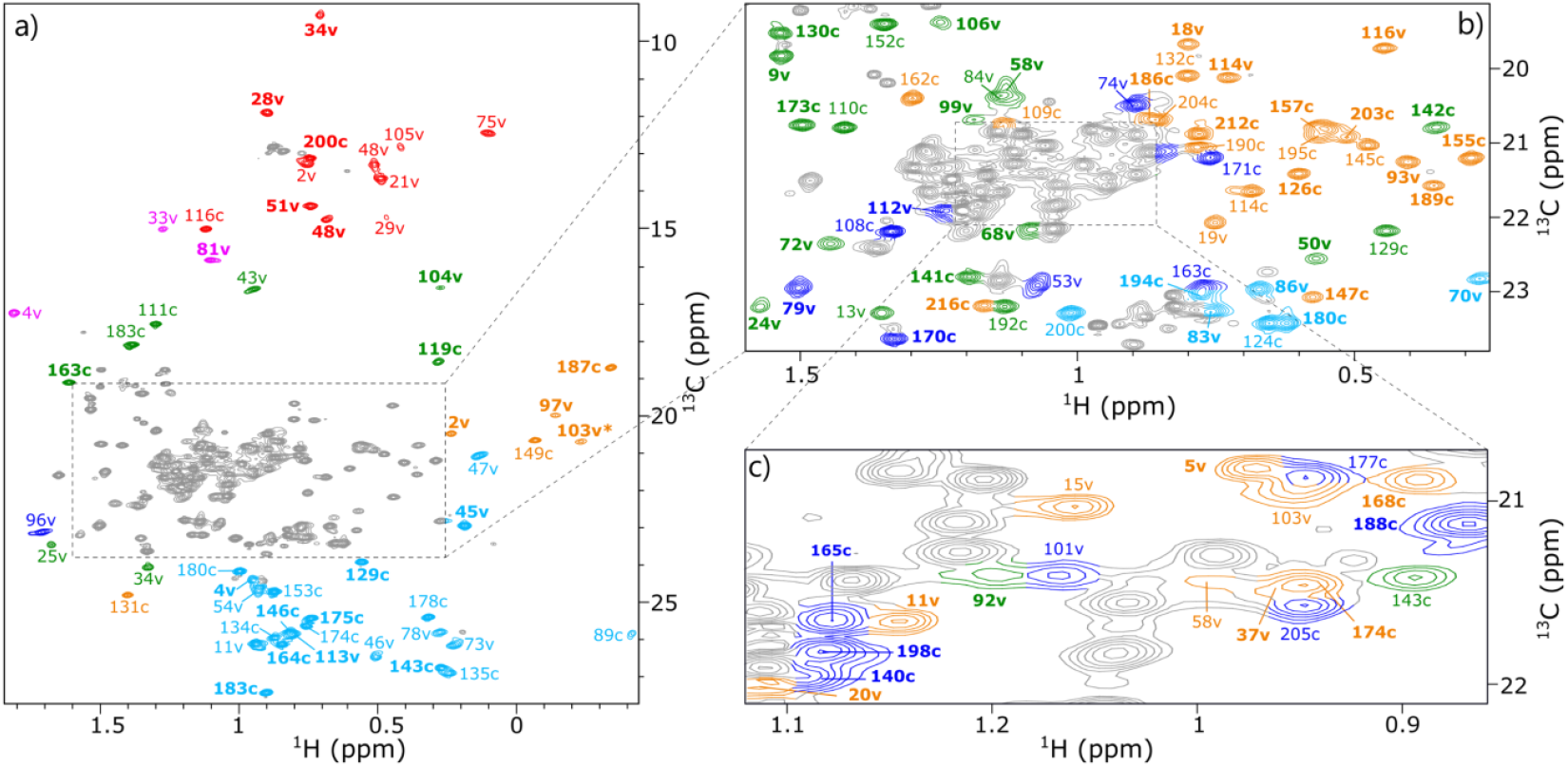
Assigned 2D ^1^H-^13^C SOFAST methyl TROSY spectrum of a Fab fragment from the antibody anti-LAMP1. The Fab fragment was produced in cell-free system and U-[^2^H], Ala-[^13^C^1^H_3_]^β^, Met-[^13^C^1^H_3_]^ε^, Leu-[^13^C^1^H_3_]^*pro- S*^, Val-[^13^C^1^H_3_]^*pro-R*^, Ile-[^13^C^1^H_3_]^δ1^, Thr-[^13^C^1^H_3_]^γ^ specifically labelled. Each assigned signal is annotated with the corresponding residue number. Alanine-β, isoleucine-δ_1_, valine-γ-*pro-R*, leucine-δ-*pro-S*, threonine-γ, and methionine-ε methyl signals are depicted in green, red, orange, light blue, dark blue, and purple, respectively. In bold are resonances belonging to the heavy chain of the Fab and in regular the ones belonging to the light chain of the Fab. Annotated with the letter “c” are residues belonging to the constant part and with the letter “v” residues belonging to the variable part. The asterisk indicates that contour level of the residue 104 has been multiplied by 3. Signals depicted in grey are either impurities, residues from the tags or unassigned signals. **a)** Full spectrum. **b)** Zoom from panel a. **c)** Zoom from panel b, contour level has been decreased by 2

Threonines were more difficult to assign as most of them are facing the solvent and either exhibited mainly intermethyl NOE cross peaks with other non-assigned threonines or showed no intermethyl NOE connectivity at all. The use of 3D “out and back” HC(C)C and HC(CC)C experiments to connect assigned backbone resonances of threonines to their methyl group was considered, but the Thr-[1,2,3-^13^C, 3-^2^H_2_, [^13^C^1^H_3_]^γ^] labelled amino acid is not commercially available. In absence of H^β^ deuteration of Thr, the low resolution in the ^1^H-dimension (Velyvis et al. 2012) precludes the transfer of assignment from the backbone for most of the overlapping solvent-exposed threonine side chains. Nonetheless, using the NOESY spectrum, 40% of the threonine-γ methyls could be assigned (Fig. 2).

Although our *in vitro* production strategy (Giraud et al. 2024) to assign methyl groups of the anti-LAMP1 Fab does not require protein refolding as the cell-free system allows the control of disulfide bond formation during synthesis, it required high-scale productions of deuterated and specifically labelled samples. These latter increased tremendously the production cost of the samples due to both expensive labelled amino acids and large volumes of cell-free reaction resulting from the inherently low production yield in D_2_O. Additionally, the time required to produce deuterated samples is impacted by the time to source the differently labelled amino acids, the preparation of the different amino acid mixes and the yield optimization. An extensive analysis of triple-resonance experiments was done to reach 84% of labelled methyl group assignment, tag excluded.

### Assignment transfer of Fab’s constant region

To speed up the assignment of therapeutic Fab fragments, a method based on the use of this first assignment of anti-LAMP1’s methyl groups and sequence identities with targeted Fabs was investigated. A Fab fragment consists of a light and a heavy chain, each composed of a highly conserved constant domain and a variable domain. The strategy aimed at transferring the assignment from the conserved domains from anti-LAMP1 to other IgG1 Fabs sharing the same constant part and assigning the variable domains using smaller constructs.

Previously, Gagné et al. and Sarker and Aubin (Gagné et al. 2024, Sarker and Aubin 2024) published assignments of trastuzumab-scFab and adalimumab-scFab isoleucine, leucine and valine methyl groups. The sequence identity is 81% between anti-LAMP1 and trastuzumab scFab fragments and 80% between anti-LAMP1 and adalimumab scFab fragments. Additionally, the constant parts of the Fab fragments showed 100% identity with anti-LAMP1 constant part. To validate our approach, we compared I-δ_1_, V-γ-*pro-R*, and L-δ-*pro-S* assignments of anti-LAMP1’s constant part with the stereospecific assignment of trastuzumab-scFab’s constant part (Gagné et al. 2024). Even if the stereospecific NMR assignment of valines and leucines was conducted differently from us (by using 10% ^13^C and 90% ^12^C as the sole carbon source), both assignments seemed to coincide (Fig. S1) with minor differences that could arise due to different experimental conditions such as the buffer pH and temperature during NMR acquisition. As both constant parts showed similar resonances, it supported our approach to transfer NMR assignments between identical constant parts from different Fab fragments, using identical experimental conditions.

Ipilimumab’s Fab fragment was selected to establish a proof of concept demonstrating the feasibility of the assignment transfer between two IgG1 Fabs. Ipilimumab is a commercial IgG1 mAb, its Fab shares 78% identity with the anti-LAMP1’s Fab and 100% identity was found between the constant parts of ipilimumab and anti-LAMP1. As already highlighted with the constant part assignments of trastuzumab and anti-LAMP1, we managed to easily transfer 98% of the assignment from the constant part of anti-LAMP1 to ipilimumab’s Fab fragment. Spectra were collected in the same experimental conditions, and, by simple superimposition, the assignment could be transferred smoothly (Fig. 3a/b, S2a/b/c, S3a/b, S4a/b). We note that under identical experimental conditions the transfer is significantly easier.

**Fig. 3.**
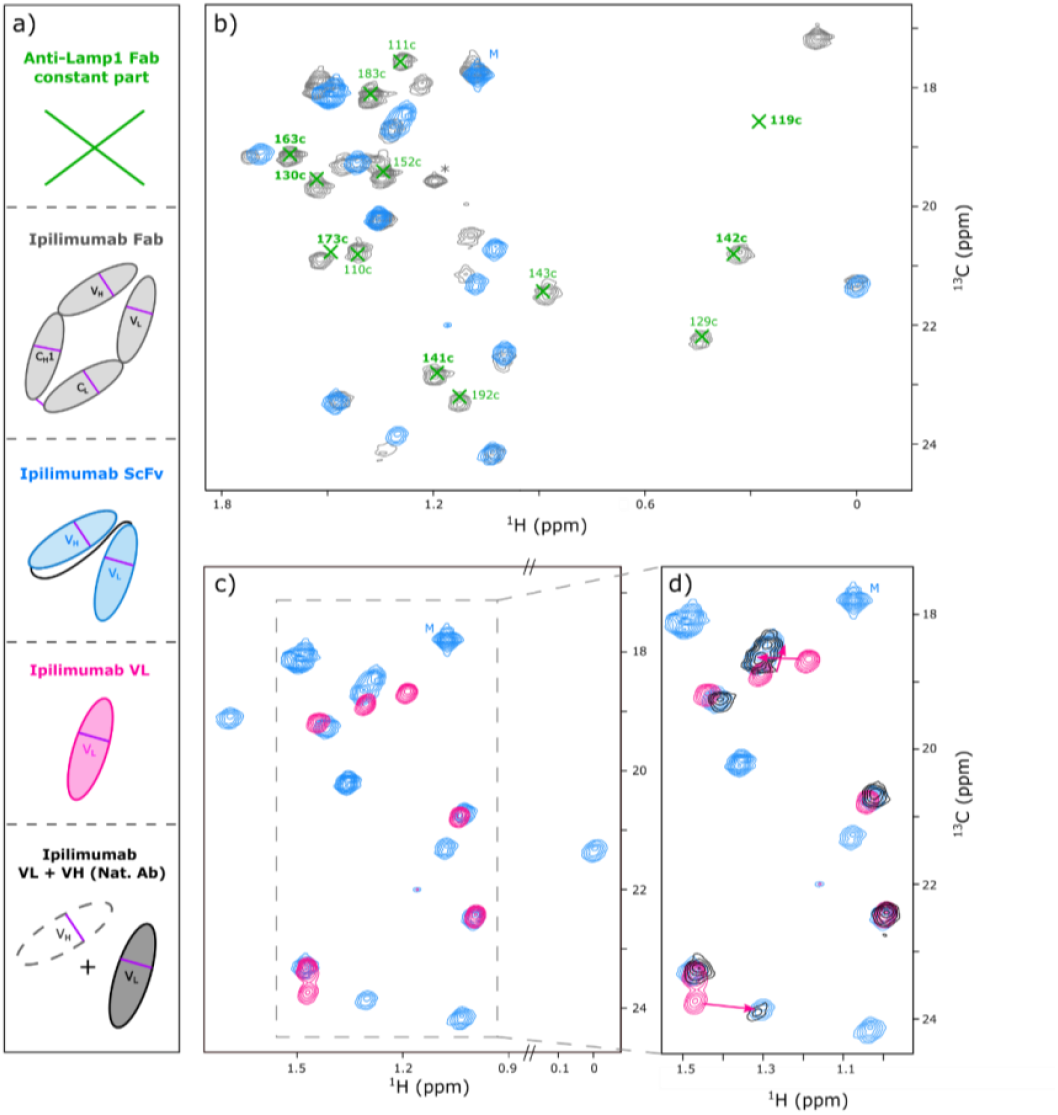
Ipilimumab assignment transfer strategy shown for the alanines. **a)** Color coded schemes of the different constructs produced and for which spectra are displayed in b, c, and d. All spectra displayed are from constructs labelled on Ala-[^13^C^1^H_3_]^β^ and Met-[^13^C^1^H_3_]^ε^. **b)** Superimposition of ^1^H-^13^C SOFAST methyl TROSY spectra of the Fab of ipilimumab in grey with the ScFv of ipilimumab in light blue. Annotated green crosses correspond to assigned signals belonging to the constant part of the Fab anti-LAMP1. In bold are resonances belonging to the heavy chain and in regular the ones belonging to the light chain. The asterisk indicates an impurity. **c)** Superimposition of ^1^H-^13^C SOFAST methyl TROSY spectra of ipilimumab’s ScFv in light blue with ipilimumab’s VL in pink. **d)** Zoom from panel c. The additional spectrum in black correspond to ipilimumab’s VL to which has been added the VH fragment in natural abundance. Arrows indicate the peaks’ shifts of the VL fragment signals upon addition of the VH fragment. A small peak doubling is observed in the ^13^C dimension for alanines’ resonances. This is due to the use of U-[^2^H], ^13^C^1^H_3_ – alanine to produce our sample in ^1^H_2_O, resulting in incorporation of ^1^H isotope in the α position due to endogenous enzymatic activities of transaminases

### A divide and conquer strategy for methyl group assignment of ipilimumab’s variable part

Two strategies were considered to assign the variable domains of ipilimumab Fab. We could study them either together as an ScFv fragment or separately using VL and VH domains. Working with the entire ScFv allows preservation of the interface between VL and VH as in the full Fab. However, the mass of ScFv is 29 kDa, and its assignment is complex without deuteration. Producing VL and VH individually gives smaller constructs that are easier to assign; however, the VH-VL interface is not preserved if the domains are produced separately, and some shifts can be observed upon transfer to the entire Fab. Addition of unlabelled VH to labelled VL could enable recovery of the interface between the two fragments. Ideally, obtaining both VL and VH domains soluble and concentrated is the most desirable option.

In our case, ipilimumab’s VL was easily produced, and its methyl group resonances were assigned. However, in our hands, the VH construct could not be obtained at concentrations enabling NMR sequential assignment. Protein precipitation occurred during sample concentration and upon buffer changes. Hence, production of the ScFv construct was also undertaken. In practice, although we have outlined both the advantages and limitations of each method, the final strategy was guided by the outcome of wet-lab production. With this divide and conquer method, the suitable size of the different constructs allowed us to produce them all in protonated buffer either uniformly ^15^N,^13^C-labelled or specifically methyl-labelled to assign each of these smaller Fab constructs.

Sequential assignments of both the light chain’s variable domain and the single-chain variable fragment of ipilimumab were performed using a combination of BEST and BEST-TROSY HNCA, HN(CA)CB, HNCO, HN(CA)CO, HN(CO)CA and HN(COCA)CB triple-resonance experiments (Favier and Brutscher 2019; Lescop et al. 2007) – see Table S3.

- VL domain

For the VL construct, 97% of ^15^N, 84% of C^α^, 84% of C’, and 60% of the C^β^ resonances were detected, excluding proline, tryptophan and cysteine residues. Indeed, the U-[^1^H, ^15^N, ^13^C] sample was produced in a cell-free system using a rich medium (hydrolyzed isogro®) that lacks free tryptophan and cysteine residues. Therefore, these latter amino acids were supplemented in the cell-free reaction, but only in their unlabelled form for cost reasons. Among the detected signals, 63% of ^15^N, 72% of C^α^, 68% of C’, and 70% of C^β^ could be assigned, tag included. Using hCCH-TOCSY experiments with mixing times of 10 ms and 20 ms, a transfer from the assigned backbone resonances to the methyl groups was undertaken. Using C^α^ and C^β^ resonances, 71% of the methyl groups of the VL construct could be assigned, tag excluded. To complete the VL assignment, a 3D HCH NOESY experiment was acquired (see Table S3). The previously assigned methyl residues, together with the observed NOE cross-peaks, allowed the assignment of the remaining five threonine and six leucine methyl groups (Fig. 4a).

**Fig. 4.**
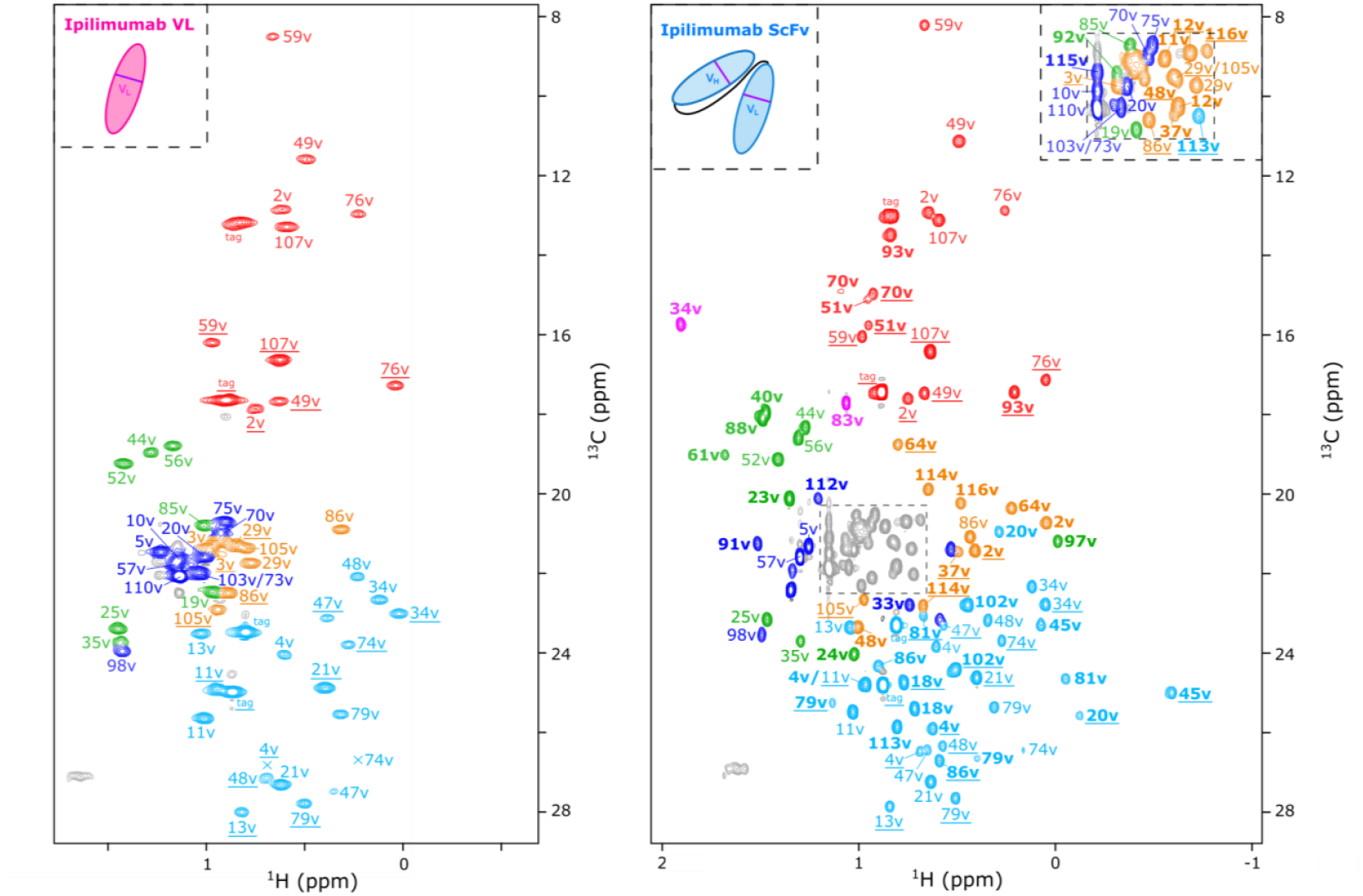
Assigned 2D ^1^H-^13^C SOFAST methyl TROSY spectra of the VL fragment of ipilimumab (left) and the ScFv of ipilimumab (right). VL and ScFv fragments were U-[^1^H, ^15^N, ^13^C] labelled. Each signal is annotated with the corresponding residue number. Alanines-β, isoleucines-δ_1_, valines-γ-*pro-R*, leucines-δ-*pro-S*, threonines-γ, and methionines-ε methyl signals are depicted in green, red, orange, light blue, dark blue, and purple, respectively. In bold are resonances belonging to the heavy chain and in regular the ones belonging to the light chain. Residues numbers are underlined for resonances of I-γ_2_, L-δ-*pro-R*, and V-γ*-pro-S*

- ScFv construct

For the ScFv construct, 92% of ^15^N, 86% of C^α^, 88% of C’, and 59% of C^β^ were detected, excluding proline, tryptophan and cysteine residues. Among the detected residues, 58% of ^15^N, 62% of C^α^, 59% of C’, and 65% of C^β^ could be assigned. Using an hCCH-TOCSY experiment together with the backbone assignment of ScFv and the already assigned methyl spectrum of the VL domain, 56 out of 74 methyl residues (76%) of ScFv were assigned, tag excluded, either by transfer from the backbone to the methyl groups or by transfer from the VL to the ScFv. All methyl groups belonging to the light chain of the ScFv were assigned, except alanine 35v and valine 3v *pro-R*. Some shifts were observed between the spectra of the ScFv and the VL (Fig. 1c, S2d, S3c, S4c), complicating the assignment transfer, but these shifts were expected for residues in very close proximity to the variable heavy chain of the ScFv. Therefore, we introduced some VH at natural abundance into our labelled VL samples and followed the shifts as we reproduced the environment found in the ScFv (Fig. 1d, S2e, S3d, S4d). This allowed us to assign alanine 35v in the ScFv construct.

To complete the assignment of methyl groups belonging to the variable heavy chain of the ScFv, the 0.22 mM sample of the U-[^1^H, ^15^N, ^13^C] ScFv ipilimumab construct was re-employed to acquire a 3D HCH NOESY experiment. Nine additional methyl residues (13%) were assigned using methyl NOEs. Two alanine residues in the variable heavy chain, 23v and 97v, initially remained unassigned. However, only alanine 97v is in close proximity to both a phenylalanine and a tryptophan, supporting the presence of a ring current effect and explaining the downfield shift of its ^1^H chemical shift. Based on this observation, both 23v and 97v could be confidently assigned and the percentage of assigned methyl groups was further improved to 92%, tag excluded (Fig. 4b). Assigning methyl group signal of a 29 kDa protein without deuteration was a significant challenge. The high percentage of assignment obtained could be further improved by detecting long-range methyl-methyl NOEs (Kerfah et al. 2015; Ayala et al. 2020). This latter would, however, require the complex large-scale production of a deuterated and methyl labelled sample.

### Methyl group assignment of ipilimumab’s Fab

By transferring the assignment from the anti-LAMP1 Fab constant part and ipilimumab’s ScFv to ipilimumab’s Fab we could transfer 91% of valine, 93% of alanine, 100% of methionine, 100% of isoleucine, 91% of threonine, and 80% of leucine residues’ assignments. As assignments were not complete for the anti-LAMP1’s constant part and ipilimumab’s variable part, the percentage of ipilimumab’s Fab assignment is lower than the percentage of assignment transfer. Nonetheless, we reached 81% of ipilimumab’s Fab assignment, tag excluded, without having to produce it deuterated (Fig. 5).

**Fig. 5.**
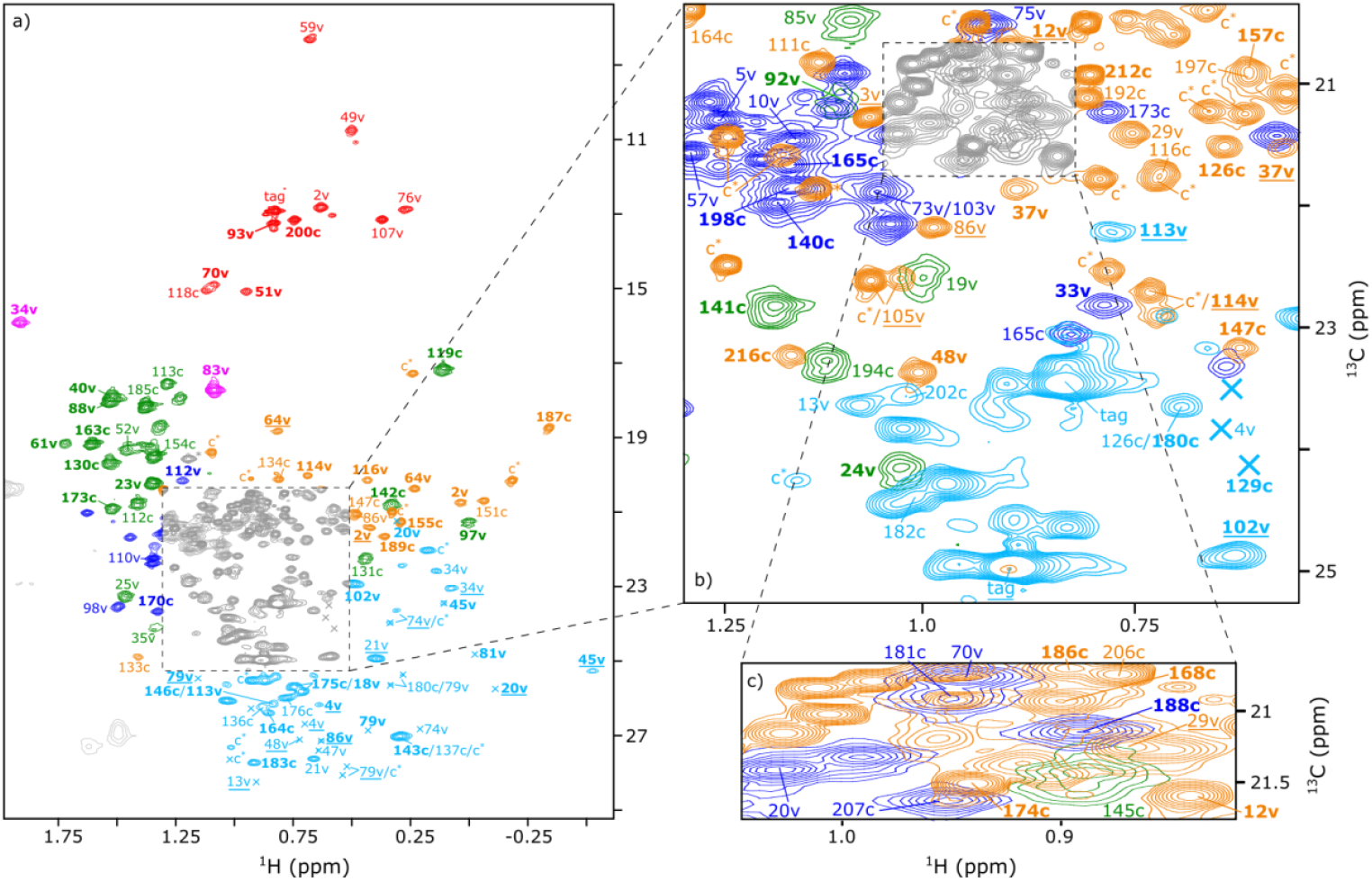
Assigned 2D ^1^H-^13^C SOFAST methyl TROSY spectrum of a Fab fragment from the antibody Ipilimumab. The spectrum is a composite figure created by superimposition of four 2D ^1^H-^13^C SOFAST methyl TROSY spectra of Ipilimumab’s Fab labelled on Met-[^13^C^1^H_3_] and Ala-[^13^C^1^H_3_]^β^; Met-[^13^C^1^H_3_] ^ε^, Ile-[^13^C^1^H_3_]^δ1^ and Thr-[^13^C^1^H_3_]^γ^; Met-[^13^C^1^H_3_]^ε^ and Val-[^13^C^1^H_3_]^*pro-S&R*^ and Met-[^13^C^1^H_3_]^ε^ and Leu-[^13^C^1^H_3_]^*pro-S&R*^. Alanines, isoleucines, valines, leucines, threonines, and methionines resonances are depicted in green, red, orange, light blue, dark blue, and purple, respectively. Each assigned signal is annotated with the corresponding residue number. The asterisk indicates an impurity. In bold are resonances belonging to the heavy chain and in regular the ones belonging to the light chain. Annotated with the letter “c” are residues belonging to the constant part and with the letter “v” residues belonging to the variable part. Residues numbers are underlined for resonances of I-γ_2_, L-δ-*pro-R*, and V-γ*-pro-S*. “c^*^” indicate resonances of leucines-δ-*pro-R* and valines-γ*-pro-S* from constant part. **a)** Full 2D spectrum. **b)** Zoom from panel a. **c)** Zoom from panel b.

The limitations of this method lie in the fact that some residues in the ScFv construct have different surroundings compared to the same residues in the Fab fragment. This could potentially induce chemical shift perturbations for residues in the variable part located at the interface with the constant part, thereby complicating the assignment transfer from the isolated variable part to the entire Fab. However, this interface between the variable and the constant parts is relatively limited and contains very few methyl residues. The variable regions of both the light and heavy chains are connected to their respective constant parts with a loop constituted of approximately 6-8 residues. In our case, five methyl groups from the variable region are located less than 6 Å away from the constant domains, and six residues from the constant domains are within 6 Å of the variable region. These eleven residues account for only a small fraction of the residues targeted for assignment transfer. As previously discussed, the assignment of the six residues belonging to the constant part and close in space to the variable part (alanine 110c, valines 109c and 162c and threonines 163c and 171c from the light chain and alanine 119c from the heavy chain) was easily transferred from the already assigned anti-LAMP1 Fab to ipilimumab (Fig. 1b, S2a/b, S4a/b). This observation could be explained by the high percentage of identity between the two Fabs and very similar sequences in the variable region close to the constant domain (Fig. S5). On the contrary, transferring the assignment of the five residues belonging to the variable part and close in space to the constant part using the ScFv is slightly more complex as some shifts appeared due to chemical environmental changes between the ScFv and the entire Fab, but they represent only 3% of the total residue number.

## Conclusion

In this paper, we developed a strategy based on the “divide and conquer” method, that aims at assigning smaller fragments and transferring their assignments to an entire Fab fragment without having to produce any deuterated sample. By combining this new method with the sequence identity of the highly conserved constant domains of Fab fragments, we successfully assigned 81% of the methyl groups of the ipilimumab Fab fragment (excluding the tag). While assigning the methyl groups of our first anti-LAMP1 Fab fragment was time-consuming, the new method enabled the assignment process for a second IgG1 Fab fragment to be completed in only one third of the time. This faster approach involved producing samples, assigning backbone and methyl groups of smaller fragments, and transferring assignments to the full Fab. This significant improvement highlights the efficiency of our approach for accelerating Fab fragment characterization. This new approach can be extended to all IgG1 Fabs provided that they share 100% identity in their constant part with anti-LAMP1. As previously discussed, success and speed of the strategy depend primarily on the nature and number of constructs that express efficiently and that are soluble. If both VL and VH domains can be individually expressed and are soluble, the NMR assignment of the variable region will be facilitated. Titration experiments of VL with VH domain (or vice versa) could be performed to complete the assignment of residues at the interface.

## Supplementary Information

The online version contains supplementary material available at XXXXXX

## Acknowledgements

The authors thank Isabel Ayala, Rida Awad, Yoan Riquier Marcussy, Nicola Salvi, Alexey Rak and Laurent Duhau for advice, support and stimulating discussions. This work is supported by the French National Research Agency in the framework of the “Investissements d’avenir” program (ANR-15-IDEX-02), NMR4mAbs project n° ANR-22-CE29-0024-01 and by CEA/CNRS/UGA/SANOFI collaborative research programs 22CUF10215/221024 and 2023-0666/C44988/231021. This work used the high field NMR and Cell-Free facilities at the Grenoble Instruct-ERIC Center (ISBG; UAR 3518 CNRS-CEA-UGA-EMBL) within the Grenoble Partnership for Structural Biology (PSB). Platform access was supported by FRISBI (ANR-10-INBS-05-02) and GRAL, a project of the University Grenoble Alpes graduate school (Ecoles Universitaires de Recherche) CBH-EUR-GS (ANR-17-EURE-0003) and IR INFRANALYTICS FR2054. IBS acknowledges integration into the Interdisciplinary Research Institute of Grenoble (IRIG, CEA).

## Author contributions

F.H., B.V., A.G., S.C., E.C., C.D., J.B., and O.F. designed the experiments. F.H., B.V., A.G., S.D., and L.I. prepared the samples. F.H., B.V., A.G., and J.B. collected the NMR data, F.H., B.V., A.F., P.G., and J.B. analyzed NMR data. F.H., B.V., J.B., and O.F. wrote the manuscript. F.H. and B.V. prepared the figures. All authors reviewed the manuscript and approved the final version.

## Data availability

chemical shifts and Bruker raw data ser files will be deposited in the BMRB data bank with entry numbers XXXXX, XXXXX, XXXXX and XXXXX.

