## Additional information for "A fast and efficient strategy for the NMR assignment of Fab methyl groups"

### Supplementary information

**Sequences** of the two chains of anti-LAMP1

**Sequences** of the five constructs from ipilimumab

**Supplementary table S1**: Optimized concentrations of magnesium, DsbC and betaine

**Supplementary table S2**: Samples and NMR experiments acquired on anti-LAMP1 Fab

**Supplementary table S3**: Samples and NMR experiments acquired on ipilimumab constructs

**Supplementary table S4**: Experimental details for the purification of ipilimumab constructs

**Figure S1:** Superimposition of anti-LAMP1 Fab with trastuzumab scFab constant part resonances

**Figure S2:** Ipilimumab assignment transfer strategy shown for the valines

**Figure S3:** Ipilimumab assignment transfer strategy shown for the isoleucines

**Figure S4:** Ipilimumab assignment transfer strategy shown for the threonines

**Figure S5:** Sequence alignments comparing anti-LAMP1 and ipilimumab's Fab fragments

**Sequences of the two chains of anti-LAMP1**

**HC**

MGWSHPQFEKGGGSGGGSGGSAWSHPQFEKENLYFQGQVQLVQSGAEVKKPGSSVKVSCKASGYIFT
NYNIHWVKKSPGQGLEWIGAIYPGNGDAPYSQKFQGKATLTADTSTSTTYMELSSLRSEDTAVYYCVRAN
WDVAFAYWGQGTLVTVSSASTKGPSVFPLAPSSKSTSGGTAALGCLVKDYFPEPVTVSWNSGALTSGVHT
FPAVLQSSGLYSLSSVVTVPSSSLGTQTYICNVNHKPSNTKVDKKVEPKSCDKTHT

**LC**

MSGSHHHHHHSSGIEGRGRENLYFQGDIQMTQSPSSLSASVGDRVTITCKASQDIDRYMAWYQDKPG
KAPRLLIHDTSTLQSGVPSRFSGSGSGRDYTLTISNLEPEDFATYYCLQYDNLWTFGGGTKVEIKRTVAAPS
VFIFPPSDEQLKSGTASVVCLLNNFYPREAKVQWKVDNALQSGNSQESVTEQDSKDSTYSLSSTLTLSKA
DYEKHKVYACEVTHQGLSSPVTKSFNRGEC

* With regards to the previously published paper from Giraud et al. (Giraud et al. 2024) the sequence numbering of anti-LAMP1 heavy and light chains has been shifted by 1 for both chains so that the sequence starts at D1 for the light chain and Q1 for the heavy chain.

**Sequences of the five constructs from ipilimumab**

**VL**

MSGSHHHHHHSSGIEGRGRENLYFQGEIVLTQSPGTLSLSPGERATLSCRASQSVGSSYLAWYQQKPG
QAPRLLIYGAFSRATGIPDRFSGSGSGTDFTLTISRLEPEDFAVYYCQQYGSSPWTFGQGTKVEIKRT

**VH**

MSGSHHHHHHSSGIEGRGRENLYFQGQVQLVESGGGVVQPGRSLRLSCAASGFTFSSYTMHWVRQA
PGKGLEWVTFISYDGNNKYYADSVKGRFTISRDNSKNTLYLQMNSLRAEDTAIYYCARTGWLGPFDYWG
QGTLVTVSS

**ScFv**

MSGSHHHHHHSSGIEGRGRENLYFQGQVQLVESGGGVVQPGRSLRLSCAASGFTFSSYTMHWVRQA
PGKGLEWVTFISYDGNNKYYADSVKGRFTISRDNSKNTLYLQMNSLRAEDTAIYYCARTGWLGPFDYWG
QGTLVTVSS**GGGGSGGGGSGGGGSGGGGS**EIVLTQSPGTLSLSPGERATLSCRASQSVGSSYLAWYQ
QKPGQAPRLLIYGAFSRATGIPDRFSGSGSGTDFTLTISRLEPEDFAVYYCQQYGSSPWTFGQGTKVEIKRT

**HC**

MSGSHHHHHHSSGIEGRGRENLYFQGQVQLVESGGGVVQPGRSLRLSCAASGFTFSSYTMHWVRQA
PGKGLEWVTFISYDGNNKYYADSVKGRFTISRDNSKNTLYLQMNSLRAEDTAIYYCARTGWLGPFDYWG
QGTLVTVSSASTKGPSVFPLAPSSKSTSGGTAALGCLVKDYFPEPVTVSWNSGALTSGVHTFPAVLQSSGL
YSLSSVVTVPSSSLGTQTYICNVNHKPSNTKVDKKVEPKSCDK

**LC**

MGEIVLTQSPGTLSLSPGERATLSCRASQSVGSSYLAWYQQKPGQAPRLLIYGAFSRATGIPDRFSGSGS
GTDFTLTISRLEPEDFAVYYCQQYGSSPWTFGQGTKVEIKRTVAAPSVFIFPPSDEQLKSGTASVVCLLNNF
YPREAKVQWKVDNALQSGNSQESVTEQDSKDSTYSLSSTLTLSKADYEKHKVYACEVTHQGLSSPVTKSF
NRGEC

**Supplementary table S1** Concentrations of magnesium, DsbC and betaine optimized for the different constructs

| | Fab anti-LAMP1 | Fab Ipilimumab | ScFv Ipilimumab | VH Ipilimumab | VL Ipilimumab |
|---|---|---|---|---|---|
| **Magnesium (mM)** | 14 | 14 | 14 | 14 | 14 |
| **DsbC (µM)** | 14 | 20 | 11 | 10 | 0 |
| **Bétaine (mM)** | 15 | 12 | 9 | 8 | 10 |

**Supplementary table S2** Summary of samples' labelling and NMR experiments acquired on anti-LAMP1 Fab for its assignment

| Sample | Labelling | Quantity | Experiment | Time | Magnetic field |
|---|---|---|---|---|---|
| Fab anti-LAMP1 | Met, Ala: $^{13}CH_3$ | 390 µg<br>180 µL at 41 µM | SOFAST HMQC | 2.25 h | 950 MHz |
| Fab anti-LAMP1 | Met, Ile-$\delta_1$: $^{13}CH_3$ | 450 µg<br>100 µL at 84 µM | SOFAST HMQC | 3.5 h | 950 MHz |
| Fab anti-LAMP1 | Met, Val-γ-*pro-R*: $^{13}CH_3$ | 440 µg<br>200 µL at 41 µM | SOFAST HMQC | 13.5 h | 950 MHz |
| Fab anti-LAMP1 | Met, Thr: $^{13}CH_3$ | 784 µg<br>220 µL at 66 µM | SOFAST HMQC | 2.75 h | 950 MHz |
| Fab anti-LAMP1 | U-[$^{2}H$]; Met, Ala, Ile-$\delta_1$, Val-γ-*pro-R*, Thr, Leu-δ-*pro-S*: $^{13}CH_3$ | 3.0 mg<br>180 µL at 314 µM | CCH HMQC-NOESY-HMQC | 12 d | 950 MHz |
| Fab anti-LAMP1 | U-[$^{2}H$]; Ala, Ile-$\delta_1$, Val-γ-*pro-R*: U-[$^{13}C$], $^{13}CH_3$ | 2.5 mg<br>180 µL at 258 µM | HCC<br>HC(C)C<br>HC(CC)C | 1 d 20 h<br>3 d 21 h<br>5 d 15 h | 950 MHz |

**Supplementary table S3** Summary of samples' labelling and NMR experiments acquired on the different ipilimumab constructs for their assignments

| Sample | Labelling | Quantity | Experiment | Time | Magnetic field |
|---|---|---|---|---|---|
| VL Ipilimumab | Met, Ala: $^{13}CH_3$ | 550 µg<br>250 µL at 140 µM | SOFAST HMQC | 1 h | 700 MHz |
| VL Ipilimumab | Met, Ile-$\delta_1$, Val-γ-*pro-S*: $^{13}CH_3$ | 340 µg<br>250 µL at 95 µM | SOFAST HMQC | 3 h | 700 MHz |
| VL Ipilimumab | Met, Thr: $^{13}CH_3$ | 380 µg<br>250 µL at 108 µM | SOFAST HMQC | 0.5 h | 850 MHz |
| VL Ipilimumab | Met, Val-γ-*pro-R*, Val-γ-*pro-S*: $^{13}CH_3$ | 105 µg<br>150 µL at 45 µM | SOFAST HMQC | 6.5 h | 600 MHz |
| VL Ipilimumab | Met, Leu-δ-*pro-S*: $^{13}CH_3$ | 35 µg<br>150 µL at 16 µM | SOFAST HMQC | 14 h | 600 MHz |
| VL Ipilimumab | U-[$^{1}H$,$^{15}N$,$^{13}C$] | 1 mg<br>180 µL at 370 µM | CT HMQC-SOFAST | 3 h | 600 MHz |
| | | | B-HSQC | 0.5 h | 600 MHz |
| | | | hCCH TOCSY 20 ms | 3 d 14 h | 600 MHz |
| | | | B-HN(COCA)CB | 2 d 11 h | 700 MHz |
| | | | hCCH TOCSY 10 ms | 1 d 4 h | 700 MHz |
| | | | BT-HN(CA)CO | 1 d 20 h | 700 MHz |
| | | | B-HNCA | 20 h | 700 MHz |

| | | | | | |
|---|---|---|---|---|---|
| | | | B-HN(CO)CA | 1 d 17 h | 600 MHz |
| | | | BT-HN(CA)CB | 1 d 17 h | 850 MHz |
| | | | B-HNCO | 16 h | 600 MHz |
| VL Ipilimumab | U-[$^{1}$H,$^{15}$N,$^{13}$C] | 3.7 mg 180 µL at 1.38 mM | NOESY HCH | 3 d | 850 MHz |
| ScFv Ipilimumab | Met, Ala: $^{13}CH_3$ | 1 mg 200 µL at 172 µM | SOFAST HMQC | 1 h | 850 MHz |
| ScFv Ipilimumab | Met, Ile-$\delta_1$, Thr: $^{13}CH_3$ | 900 µg 200 µL at 153 µM | SOFAST HMQC | 2 h | 850 MHz |
| ScFv Ipilimumab | Met, Val-γ-*pro-R*, Val-γ-*pro-S*: $^{13}CH_3$ | 240 µg 180 µL at 47 µM | SOFAST HMQC | 3 h 30 | 600 MHz |
| ScFv Ipilimumab | Met, Ile-$\delta_1$, Val-γ-*pro-S*: $^{13}CH_3$ | 90 µg 150 µL at 20 µM | SOFAST HMQC | 14 h | 600 MHz |
| ScFv Ipilimumab | Met, Leu-δ-*pro-R*: $^{13}CH_3$ | 495 µg 150 µL at 114 µM | SOFAST HMQC | 1.2 h | 800 MHz |
| ScFv Ipilimumab | U-[$^{1}$H,$^{15}$N,$^{13}$C] | 1.5 mg 240 µL at 218 µM | CT HMQC-SOFAST | 1 h 20 | 600 MHz |
| | | | B-TROSY | 1 h 10 | 600 MHz |
| | | | CT HMQC-SOFAST | 0.67 h | 850 MHz |
| | | | B-TROSY | 1 h | 850 MHz |
| | | | BT-HN(CO)CA | 2 d 8 h | 600 MHz |
| | | | hCCH TOCSY 10ms | 3 d 9 h | 600 MHz |
| | | | BT-HNCO | 17 h | 600 MHz |
| | | | BT-HN(CA)CB | 2 d 19 h | 850 MHz |
| | | | BT-HNCA | 1d 15 h | 850 MHz |
| | | | BT-HN(CA)CO | 2 d | 850 MHz |
| | | | NOESY HCH | 3d 15 h | 800 MHz |
| Fab Ipilimumab | Met, Ala: $^{13}CH_3$ | 220 µg 200 µL at 22 µM | SOFAST HMQC | 18 h | 700 MHz |
| Fab Ipilimumab | Met, Ile-$\delta_1$, Thr: $^{13}CH_3$ | 200 µg 195 µL at 20 µM | SOFAST HMQC | 20 h | 850 MHz |
| Fab Ipilimumab | Met, Val-γ-*pro-R*, Val-γ-*pro-S*: $^{13}CH_3$ | 520 µg 180 µL at 57 µM | SOFAST HMQC | 3.5 | 950 MHz |
| Fab Ipilimumab | Met, Ile-$\delta_1$, Val-γ-*pro-S*: $^{13}CH_3$ | 270 µg 180 µL at 30 µM | SOFAST HMQC | 13 h | 600 MHz |
| Fab Ipilimumab | Met, Leu-δ-*pro-R*: $^{13}CH_3$ | 350 µg 140 µL at 128 µM | SOFAST HMQC | 14.9 h | 800 MHz |
| Fab Ipilimumab | Met, Leu-δ-*pro-R*, Leu-δ-*pro-S*: $^{13}CH_3$ | 830 µg 150 µL at 110 µM | CT SOFAST HMQC | 7.5 h | 700 MHz |
| | | | SOFAST HMQC | 4 h | |
| **Ipilimumab VL + VH** | VL Met, Ala: $^{13}CH_3$ + VH Nat. Ab. | 73 µg VL + 160 µg VH | SOFAST HMQC | 5.5 h | 600 MHz |
| **Ipilimumab VL + VH** | VL Met, Ile-$\delta_1$, Val-γ-*pro-S*: $^{13}CH_3$ + VH Nat. Ab. | 50 µg VL + 120 µg VH | SOFAST HMQC | 15 h | 600 MHz |
| **Ipilimumab VL + VH** | VL Met, Thr: $^{13}CH_3$+ VH Nat. Ab. | 74 µg VL + 160 µg VH | SOFAST HMQC | 11.5 h | 600 MHz |

**Supplementary table S4** Experimental details for the affinity chromatography purification of ipilimumab constructs. CV: column volumes

| | **HiTrap protein L 1 mL** | **HisTrap FF 1 mL** |
|---|---|---|
| **Device** | NGC chromatography system (Bio-Rad) | |
| **Flow rate** | 1 mL/min | 1 mL/min |
| **Buffer A** | 20 mM NaP, 150 mM NaCl, pH 7.4 | PBS 2X, pH 7.4 |
| **Buffer B** | 20 mM NaP, pH 7.4 | PBS 2X, 50 mM imidazole pH 7.4 |
| **Buffer C** | 0.1 M Glycine pH 3 | / |
| **Equilibration** | 5 CV buffer A | 5 CV buffer A + 3% buffer B |
| **Sample loading** | Sample diluted by 4 in buffer A and loaded at 0.6 mL/min | Sample diluted by 4 in buffer A + 3% buffer B and loaded at 0.6 mL/min |
| **Wash** | 5 CV buffer A,<br>5 CV buffer B | 8 CV buffer A + 3% buffer B,<br>8 CV buffer A + 7% buffer B,<br>8 CV buffer A + 12% buffer B |
| **Elution** | 5 CV 0-100% buffer C in buffer B,<br>5 CV buffer C<br>Fractions of 1 mL dropping in wells filled with 50 µL 1 M Tris pH 8 | 9 CV 12-100 % buffer B in buffer A, 5 CV buffer B<br>Fractions of 1 mL |

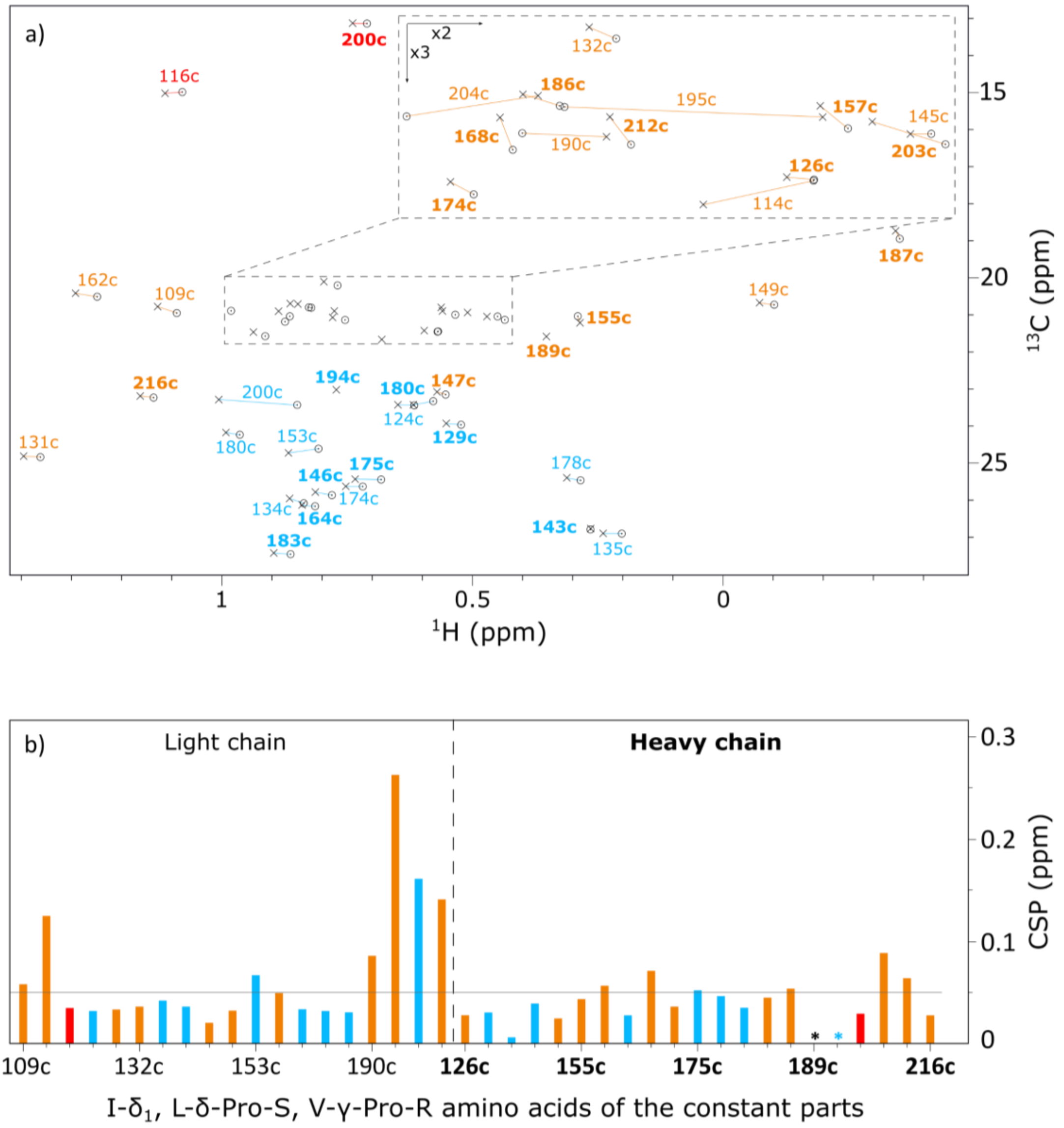


**Fig. S1** a) Superimposition of anti-LAMP1 Fab constant part resonances represented by crosses with trastuzumab scFab's ones (Gagné et al. 2024, BMRB: 52228) represented by circles. Annotated in orange, blue, and red are signals of valines-γ-*pro-R*, leucines-δ-*pro-S*, and isoleucines-$\delta_1$, respectively. In bold are resonances belonging to the heavy chain and in regular the ones belonging to the light chain. Lines are drawn to connect circles and crosses that were assigned to the same amino acid. Residues valine 189c and leucine 194c from the heavy chain were not assigned for trastuzumab-scFab. b) Chemical shift differences measured between the assignment of anti-LAMP1 Fab constant part and trastuzumab-scFab's one for valines-γ-*pro-R*, leucines-*δ-pro-S* and isoleucines-$\delta_1$. CSP were calculated using the following formula: $CSP = \sqrt{\Delta\delta_H{}^2 + \left(\frac{\Delta\delta_C}{4}\right)^2}$

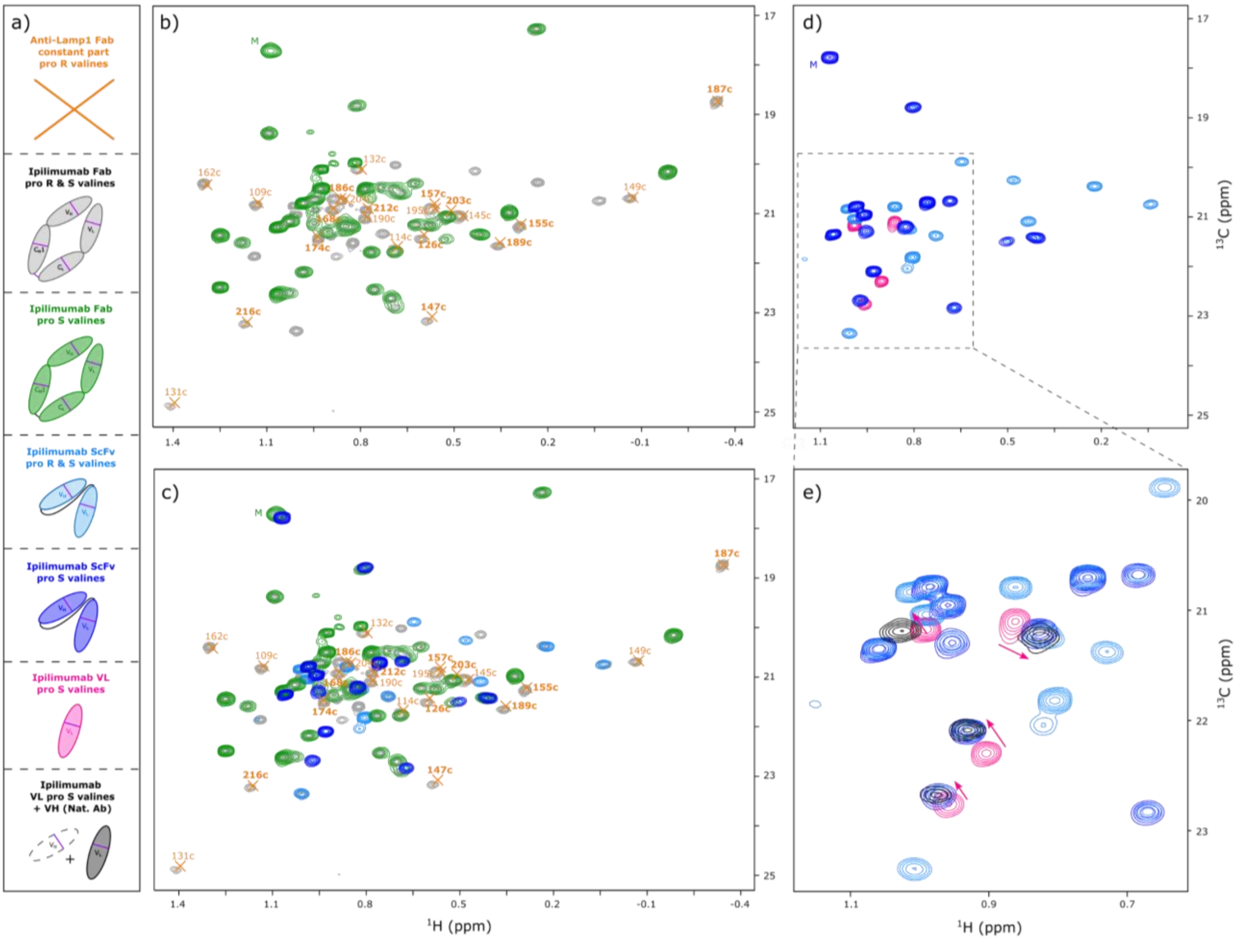


**Fig. S2** Ipilimumab assignment transfer strategy shown for the valines. a) Color coded schemes of the different constructs produced and for which spectra are displayed in b, c, d and e. All spectra displayed are from constructs labelled either on Val-[$^{13}C^{1}H_3$]$^{pro\text{-}S}$ and Met-[$^{13}C^{1}H_3$]$^{\varepsilon}$ or on Val-[$^{13}C^{1}H_3$]$^{pro\text{-}S\&R}$ and Met-[$^{13}C^{1}H_3$]$^{\varepsilon}$. b) Superimposition of $^{1}$H-$^{13}$C SOFAST methyl TROSY spectra of the Fab of ipilimumab labelled on Val-[$^{13}C^{1}H_3$]$^{pro\text{-}S\&R}$ and Met-[$^{13}C^{1}H_3$]$^{\varepsilon}$ in grey with the Fab of ipilimumab labelled on Val-[$^{13}C^{1}H_3$]$^{pro\text{-}S}$ and Met-[$^{13}C^{1}H_3$]$^{\varepsilon}$ in green. Annotated orange crosses correspond to assigned signals belonging to the constant part of the Fab of anti-LAMP1. In bold are resonances belonging to the heavy chain and in regular the ones belonging to the light chain. c) Panel b to which has been added two additional spectra. $^{1}$H-$^{13}$C SOFAST methyl TROSY spectra of ipilimumab's ScFv labelled on Val-[$^{13}C^{1}H_3$]$^{pro\text{-}S\&R}$ and Met-[$^{13}C^{1}H_3$]$^{\varepsilon}$ and of ipilimumab's ScFv labelled on Val-[$^{13}C^{1}H_3$]$^{pro\text{-}S}$ and Met-[$^{13}C^{1}H_3$]$^{\varepsilon}$ are represented in light blue and dark blue, respectively. d) Superimposition of $^{1}$H-$^{13}$C SOFAST methyl TROSY spectra of ipilimumab's ScFv labelled on Val-[$^{13}C^{1}H_3$]$^{pro\text{-}S\&R}$ and Met-[$^{13}C^{1}H_3$]$^{\varepsilon}$ in light blue with ipilimumab's ScFv labelled on Val-[$^{13}C^{1}H_3$]$^{pro\text{-}S}$ and Met-[$^{13}C^{1}H_3$]$^{\varepsilon}$ in dark blue and with ipilimumab's VL in pink. e) Zoom from panel d, an additional spectrum in black corresponding to ipilimumab's VL to which has been added the VH fragment in natural abundance is presented. Pink arrows indicate the peaks' shifts of the VL fragment signals upon addition of the VH fragment

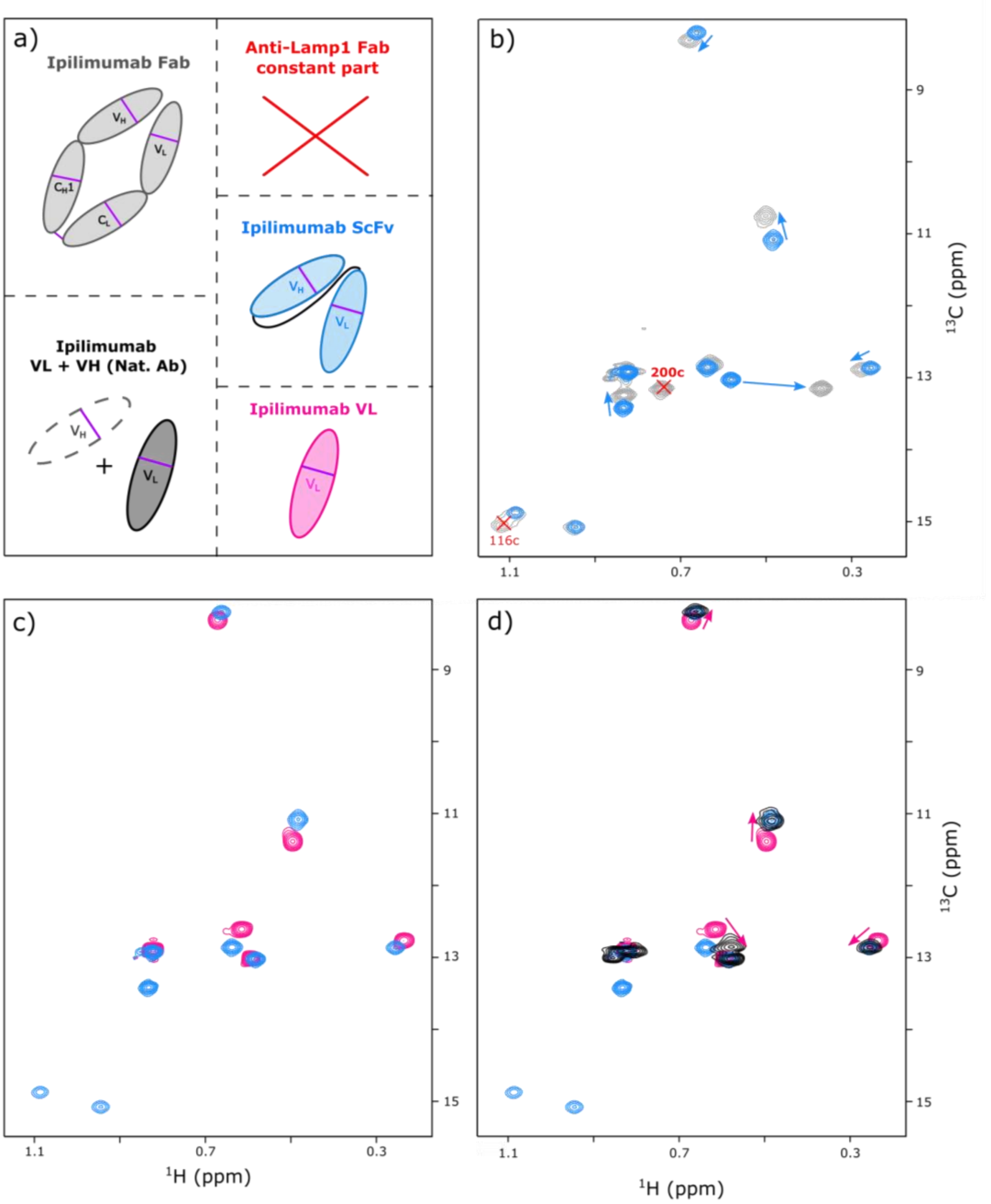


**Fig. S3** Ipilimumab assignment transfer strategy shown for the isoleucines. a) Color coded schemes of the different constructs produced and for which spectra are displayed in b, c, and d. All spectra displayed are from constructs labelled on Ile-[$^{13}C^{1}H_3$]$^{\delta 1}$and Met-[$^{13}C^{1}H_3$]$^{\varepsilon}$. b) Superimposition of $^{1}H$-$^{13}C$ SOFAST methyl TROSY spectra of the Fab ipilimumab in grey with the ScFv of ipilimumab in light blue. Light blue arrows indicate the peaks' shifts between the ScFv and the Fab fragment. Annotated red crosses correspond to assigned signals belonging to the constant part of the Fab of anti-LAMP1. In bold are resonances belonging to the heavy chain and in regular the ones belonging to the light chain. c) Superimposition of $^{1}H$-$^{13}C$ SOFAST methyl TROSY spectra of ipilimumab's ScFv in light blue with ipilimumab's VL in pink. d) Panel c, on which an additional spectrum in black corresponding to ipilimumab's VL to which has been added the VH fragment in natural abundance has been added. Pink arrows indicate the peaks' shifts of the VL fragment signals upon addition of the VH fragment

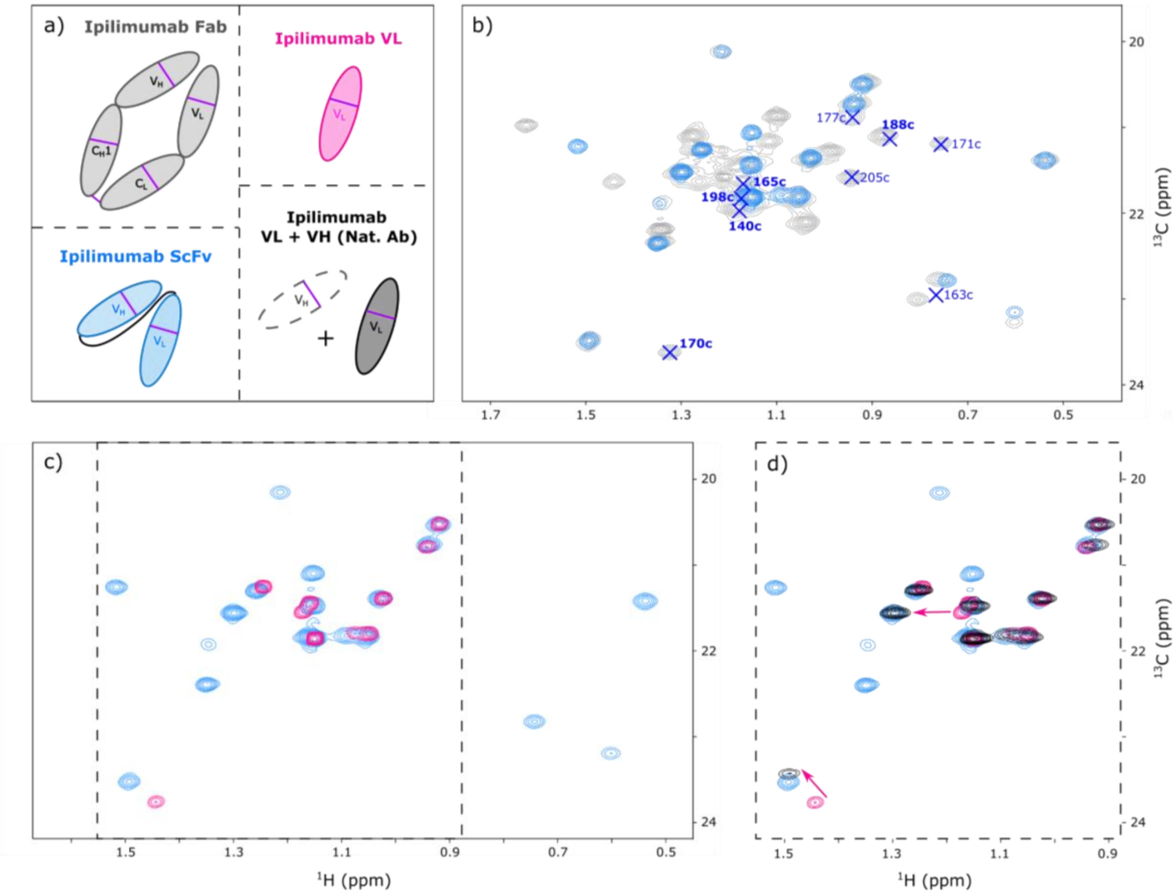


**Fig. S4** Ipilimumab assignment transfer strategy shown for the threonines. a) Color coded schemes of the different constructs produced and for which spectra are displayed in b, c, and d. All spectra displayed are from constructs labelled on Thr-[$^{13}C^{1}H_3$]γ and Met-[$^{13}C^{1}H_3$] $^{\varepsilon}$. b) Superimposition of $^{1}H$-$^{13}C$ SOFAST methyl TROSY spectra of the Fab ipilimumab in grey with the ScFv of ipilimumab in light blue. Annotated blue crosses correspond to assigned signals belonging to the constant part of the Fab of anti-LAMP1. In bold are resonances belonging to the heavy chain and in regular the ones belonging to the light chain. c) Superimposition of $^{1}H$-$^{13}C$ SOFAST methyl TROSY spectra of ipilimumab's ScFv in light blue with ipilimumab's VL in pink. d) Zoom from panel c. The additional spectrum in black corresponds to ipilimumab's VL to which has been added the VH fragment in natural abundance. Arrows indicate the peaks' shifts of the VL fragment signals upon addition of the VH fragment

**Light chain**

```
AntiLamp1  1    DIQMTQSPSSLSASVGDRVTITCKASQDI-DRYMAWYQDKPGKAPRLLIHDTSTLQSGVP  59
Ipilimumab 1    E.VL....GT..L.P.E.A.LS.R...SVGSS.L....Q...Q......YGAFSRAT.I.  60

AntiLamp1  60   SRFSGSGSGRDYTLTISNLEPEDFATYYCLQY-DNLWTFGGGTKVEIKRTVAAPSVFIFP  118
Ipilimumab 61   D........T.F.....R.......V...Q..GSSP....Q...................  120

AntiLamp1  119  PSDEQLKSGTASVVCLLNNFYPREAKVQWKVDNALQSGNSQESVTEQDSKDSTYSLSSTL  178
Ipilimumab 121  ............................................................  180

AntiLamp1  179  TLSKADYEKHKVYACEVTHQGLSSPVTKSFNRGEC  213
Ipilimumab 181  ...................................  215
```

**Heavy chain**

```
AntiLamp1  1    QVQLVQSGAEVKKPGSSVKVSCKASGYIFTNYNIHWVKKSPGQGLEWIGAI-YPGNGDAP  59
Ipilimumab 1    .....E..GG.VQ..R.LRL..A...FT.SS.TM...RQA..K....VTF.S.D..-NKY  59

AntiLamp1  60   YSQKFQGKATLTADTSTSTTYMELSSLRSEDTAVYYCVRANWDVAFAYWGQGTLVTVSSA  119
Ipilimumab 60   .ADSVK.RF.ISR.N.KN.L.LQMN...A....I...A.TG.LGP.D.............  119

AntiLamp1  120  STKGPSVFPLAPSSKSTSGGTAALGCLVKDYFPEPVTVSWNSGALTSGVHTFPAVLQSSG  179
Ipilimumab 120  ............................................................  179

AntiLamp1  180  LYSLSSVVTVPSSSLGTQTYICNVNHKPSNTKVDKKVEPKSC  221
Ipilimumab 180  ..........................................  221
```

**Fig. S5** Sequence alignments comparing the sequences of the light chains and the heavy chains of anti-LAMP1 and ipilimumab's Fab fragments. Dots represent amino acids that are identical in both sequences. Underlined amino acids are belonging to the constant parts of the Fab fragments
